# HER2 expression defines unique requirements for flotillin and c-Src for EGFR signaling

**DOI:** 10.1101/2022.04.14.488353

**Authors:** John Abousawan, Laura A. Orofiamma, Gregory D. Fairn, Costin N. Antonescu

## Abstract

The epidermal growth factor receptor (EGFR) controls many cellular functions. Upon binding its ligand, the receptor undergoes dimerization, phosphorylation, and activation of signals including the phosphatidylinositol-3-kinase (PI3K)-Akt pathway. While some studies indicated that EGFR signaling may be controlled by signal enrichment within membrane raft nanodomains, others have found a limited effect of membrane raft disruption on EGFR signaling, suggesting that specific factor(s) may define context-specific control of EGFR signaling by membrane rafts. Ligand-bound EGFR can homodimerize, or instead undergo heterodimerization with the related receptor HER2 when the latter is expressed. We examined how EGFR signaling in the presence of HER2 distinctly requires membrane raft nanodomains. Induction of HER2 expression altered EGFR signaling duration consistent with EGFR/HER2 heterodimer formation. EGFR and c-Src localized within plasma membrane structures demarked by flotillin, a membrane raft protein, selectively in HER2-expressing cells. Consistently, HER2-expressing cells, but not cells lacking HER2, were dependent on flotillin and c-Src for EGFR signaling leading to Akt activation and cell proliferation. Hence, HER2 expression establishes the requirement of EGFR signaling for flotillin membrane rafts and c-Src, leading to Akt activation.

**Summary statement:** The role of membrane rafts in EGFR signaling may be context-specific. We find that the related receptor HER2 defines unique signaling requirements for EGFR for membrane rafts, flotillin, and c-Src.

## Introduction

The epidermal growth factor (EGF) receptor (EGFR) is one of four members of the ErbB family of receptor tyrosine kinases that includes ErbB2 (HER2), ErbB3 (HER3), and ErbB4 (HER4) (Lemmon and Schlessinger, 2010). EGFR and other ErbBs play important roles in the development and homeostasis of various adult tissues, and ErbBs are also central drivers of tumor progression in a wide range of cancers (Sigismund et al., 2018), including breast cancers (Ali and Wendt, 2017; Nakai et al., 2016). Specifically, HER2 drives tumor progression in so-called HER2-positive breast cancers, while triple negative breast cancers lack HER2 expression, instead often relying on EGFR (Cancer Genome Atlas Network, 2012; Hsu and Hung, 2016; O’Reilly et al., 2015; Oh and Bang, 2019). Hence, resolving the mechanisms that regulate EGFR, HER2, and other ErbBs is thus important to better understand how these signals can drive tumor progression.

Following binding to growth factor ligands such as EGF, EGFR undergoes asymmetric receptor homodimerization, resulting in activation of its intrinsic kinase domain that in turn triggers trans-autophosphorylation of the carboxy-terminal domain of the receptor (Jura et al., 2009; Zhang et al., 2006). These phosphorylated tyrosine residues contribute to the formation of docking sites for effector proteins that contain PTB or SH2 domain such as Grb2 and Shc, resulting in the activation of intracellular signaling pathways such as phosphatidylinositol-3-kinase (PI3K)-Akt, Ras-Erk, signal transducer and activator of transcription (STAT), and phospholipase C γ1 (PLCγ1) (Wee and Wang, 2017).

Initial EGFR signaling occurs upon ligand-binding at the plasma membrane. The plasma membrane contains several distinct types of dynamic nanodomains thought to contribute to receptor signaling through scaffolding or spatiotemporal organization of signaling intermediates (Delos Santos et al., 2015; Lu and Fairn, 2018). Upon binding to ligand and concomitant to signal activation, EGFR is recruited to clathrin-coated pits (CCPs), which are 50-200 nm assemblies of clathrin, AP2 and other proteins on the inner leaflet of the plasma membrane, leading to membrane curvature and eventually EGFR internalization into vesicles (McMahon and Boucrot, 2011; Mettlen et al., 2018). Several studies showed that CCPs function not only in receptor endocytosis, but also have key roles in the regulation of specific aspects of EGFR signaling leading to activation of Akt (Cabral-Dias et al., 2022; Delos Santos et al., 2017; Garay et al., 2015; Pascolutti et al., 2019; Reis et al., 2015; Rosselli-Murai et al., 2018; Sigismund et al., 2008).This indicates that plasma membrane nanodomains such as CCPs form scaffolds essential for activation of specific intracellular signaling events.

In addition to CCPs, other structures such as membrane “rafts” may control signaling by receptors including EGFR. Membrane rafts are dynamic membrane nanodomains that are enriched in and highly dependent on cholesterol and glycosphingolipids (Lu and Fairn, 2018; Mukherjee and Maxfield, 2004; Sonnino and Prinetti, 2013). Membrane rafts are a heterogeneous ensemble of nanodomains, at least some of which are driven by specific lipid-binding proteins. Flotillin-1 and -2 are highly similar palmitoylated proteins that oligomerize and bind cholesterol, thus supporting the formation of ∼100 nm protein-driven membrane rafts that scaffold and enrich a number of receptor tyrosine kinases including EGFR and select signaling molecules (Kurrle et al., 2012; Meister and Tikkanen, 2014; Staubach and Hanisch, 2011).

There are conflicting results about the requirement for cholesterol and/or flotillin proteins and thus the involvement of membrane rafts in EGFR signaling. EGFR has been observed in membrane fractions consistent with membrane rafts, and perturbations of cholesterol alters membrane mobility of EGFR (Bag et al., 2015; Hofman et al., 2008; Irwin et al., 2011; Michio Hiroshima et al., 2021; Orr et al., 2005; Pike et al., 2005; Puri et al., 2005; Wang et al., 2009). Some studies reported that loss of membrane rafts by cholesterol disruption increases ligand affinity to EGF and increases EGFR phosphorylation and/or signaling (Chen and Resh, 2002; Lambert et al., 2008; Li et al., 2006; Liu et al., 2009; Michio Hiroshima et al., 2021; Pike and Casey, 2002; Ringerike et al., 2002; Roepstorff et al., 2002; Turk et al., 2012; Westover et al., 2003; Zhang et al., 2016). In contrast, others reported that disruption of cholesterol has no effect on EGFR signaling (Sigismund et al., 2008; Ushio-Fukai et al., 2001; Zhang et al., 2016), or instead leads to loss of EGFR signaling (Gibson et al., 2009; Hea et al., 2007; Irwin et al., 2011). Consistent with the latter, disruption of flotillin-1 leads to a loss of EGF-stimulated EGFR clustering and phosphorylation, as well as disruption of downstream signaling (Amaddii et al., 2012).

Considering that perturbations of cholesterol, flotillins, and more generally membrane rafts have inconsistent effects as reported in different studies indicates that some yet unidentified cellular factor(s) or context may define the need for membrane rafts for EGFR signaling. While EGF is specific to binding EGFR, in the presence HER2, EGF stimulation strongly favors formation of EGFR/HER2 heterodimer compared to the formation of EGFR homodimers that occurs in the absence of HER2 expression (Li et al., 2012b). While EGFR/HER2 heterodimers also lead to activation of receptor intrinsic kinase activity and receptor phosphorylation, EGFR/HER2 heterodimers exhibit differences in the extent of activation of specific signals (Hartman et al., 2012; Wolf-Yadlin et al., 2006), and in some cases exhibit delayed internalization compared to EGFR homodimers (reviewed by (Bertelsen and Stang, 2014)). Hence, co-expression of HER2 may underlie unique context-specific properties of EGFR signaling.

We previously uncovered that co-expression of HER2 rendered EGF-stimulated activation of Akt insensitive to clathrin perturbation, while clathrin was an essential regulator of EGFR signaling in cells that lack HER2 expression (Garay et al., 2015). This suggests that HER2 co-expression alters the EGFR requirement for specific clathrin-scaffolded nanodomains; however, it is not known if HER2 might influence EGFR signaling dependence on another subtype of membrane nanodomain. Importantly, HER2 can interact with flotillin-2 (Asp et al., 2014; Pust et al., 2013). While flotillin-2 can regulate HER2 stability at the plasma membrane, flotillin-1 and flotillin-2 have at least partly non-redundant functions at the plasma membrane (Bitsikas et al., 2014; Kurrle et al., 2012; Otto and Nichols, 2011), although flotillin-1 and flotillin-2 are largely present in the same plasma membrane structures (Fekri et al., 2019). This suggests that the HER2-flotillin interaction may more broadly control EGFR signaling. How flotillins and flotillin membrane rafts may contribute to the regulation of EGFR and/or HER2 signaling and thus underlie context-specific roles of membrane rafts in EGFR signaling are poorly understood.

The possibility of context-specific dependence of EGFR signaling on membrane rafts also suggests context-specific recruitment to and functional requirement for specific signaling intermediates. Unlike EGFR, HER2 can interact directly with c-Src (Muthuswamy and Muller, 1995) via its catalytic kinase domain, leading to c-Src activation and potentiation of signaling elicited by the receptor (Kim et al., 2005; Marcotte et al., 2009). Hence, HER2 co-expression may also define unique requirements for specific signaling intermediates, in addition to defining specific requirements for membrane rafts and/or other plasma membrane nanodomains.

Here, we examined how HER2 co-expression may underlie the context-specific nanoscale organization of EGFR signaling at the plasma membrane. To do so, we used MDA-MB-231 triple negative breast cancer cells and ARPE-19 cells, both of which endogenously express EGFR but not HER2 (Delos Santos et al., 2017; Garay et al., 2015). We engineered derivatives of these cell lines that co-express HER2 to allow direct comparison of signaling by EGFR homodimers and EGFR/HER2 heterodimers in a common cell background. We find that HER2 co-expression defines a functional requirement of EGFR phosphorylation and cell proliferation for membrane cholesterol, flotillin-1 and c-Src, as well as unique recruitment of EGFR and c-Src to plasma membrane flotillin nanodomains. This work resolves important context-specific requirements for flotillin membrane rafts in EGFR signaling.

## Results

### HER2 co-expression alters EGFR signaling duration

To directly compare the signaling of EGFR heterodimers and EGFR/HER2 heterodimers, we first generated a derivative cell line of MDA-MB-231 cells stably expressing HER2. While we could not detect HER2 in parental MDA-MB-231 cells, we could readily detect HER2 in MDA-MB-231-HER2 cells by western blotting and immunofluorescence staining (**Figure 1A**). The latter shows expression of HER2 in a majority of MDA-MB-231-HER2 cells.

**Figure 1.**
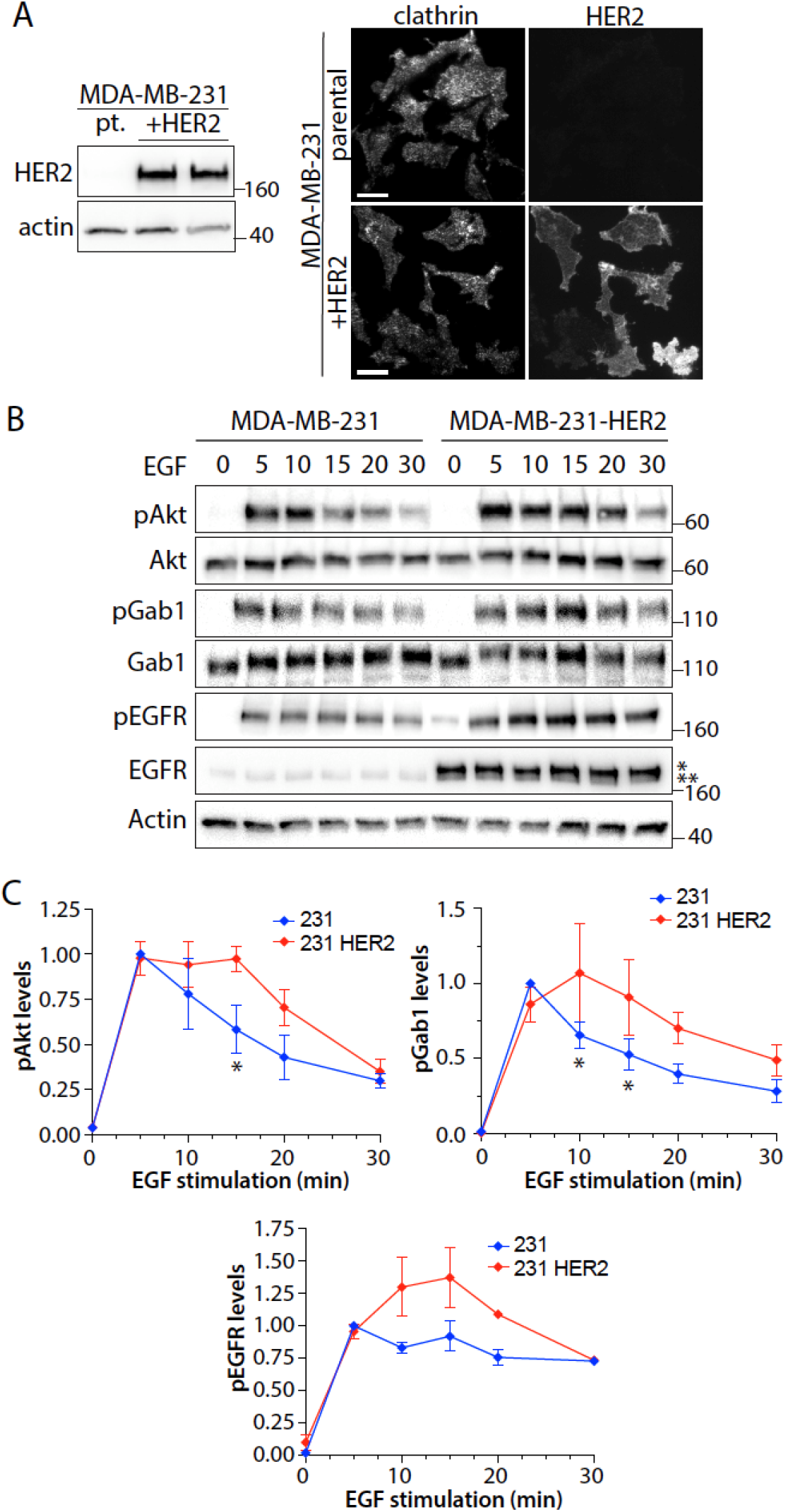
HER2 expression alters duration of EGF-stimulated signaling. (**A**) HER2 levels were detected in MDA-MB-231 (parental; pt) and MDA-MB-231-HER2 (HER2+) cells via western blotting of whole cell lysates by probing for total levels of HER2 (*left panel*). Each cell line was also subject to immunofluorescence staining of HER2. Shown (*right panel*) are representative fluorescence images obtained by TIRF microscopy, scale 40 µm. (**B**) MDA-MB-231 and MDA-MB-231-HER2 cells were stimulated with 5 ng/mL for the indicated times. Shown are representative immunoblots of whole cell lysates to detect phosphorylated proteins as indicated: pAkt (S473), pGab1 (Y627), pEGFR (Y1068) or other proteins. * and ** represent HER2 and EGFR, respectively, as detected by the anti-EGFR antibody. (**C**) Shown are the measurements of the levels of signals detected, as means ± SE normalized to 5 min EGF condition, for pAkt (s473), pGab1 (Y627), pEGFR (Y1068). All western blot panels show approximate molecular weight in kDa.

We next compared the signaling of MDA-MB-231 parental and HER2-expressing cells. Stimulation with EGF triggered robust increase in the phosphorylation of EGFR (pY1068), Gab1 (pY627) and Akt (pS473) in both cell types (**Figure 1B-C**). Notably, the antibody used to detect EGFR also detected HER2 (**Figure 1B**), and the higher expression of the higher molecular weight HER2 compared to endogenous EGFR indicates that there is abundant HER2 expression to favor EGFR/HER2 heterodimer formation over that of EGFR homodimers. Interestingly, EGF-stimulation of all three signals examined persisted longer in MDA-MB-231-HER2 cells than in MDA-MB-231 parental cells. This suggests that differences in signal activation and regulation occur at the level of EGFR phosphorylation in the context of EGFR homodimers and EGFR/HER2 heterodimers, resulting in changes in the duration of signaling following ligand stimulation. These differences in signaling by EGFR homodimers and EGFR/HER2 heterodimers are consistent with previous reports (Hartman et al., 2012; Wolf-Yadlin et al., 2006) and may reflect changes in the nanoscale organization of receptor signaling at the plasma membrane.

### Flotillin-dependent membrane rafts are required for EGFR signaling upon HER2 expression

To resolve how differences in EGFR signaling upon co-expression of HER2 may reflect distinct functional requirements for membrane raft nanodomains in EGFR homodimer and EGFR/HER2 heterodimer signaling, we examined the effect of cholesterol perturbation on EGFR signaling using nystatin treatment, a more selective agent for this purpose than methyl-β-cyclodextrin and others (Sigismund et al., 2008). Importantly, nystatin treatment led to a reduction in EGF-stimulated phosphorylation of Akt in MDA-MB-231-HER2 cells (**Figure 2 A,C**) but was without effect in parental MDA-MB-231 cells (**Figure 2A-B**). Nystatin treatment was without effect on EGF-stimulated EGFR phosphorylation in either cell line (**Figure 2 A-C**), indicating that the loss of EGF-stimulated Akt phosphorylation in MDA-MB-231-HER2 cells upon treatment with nystatin was not due to impaired ligand-binding to cell surface EGFR.

**Figure 2.**
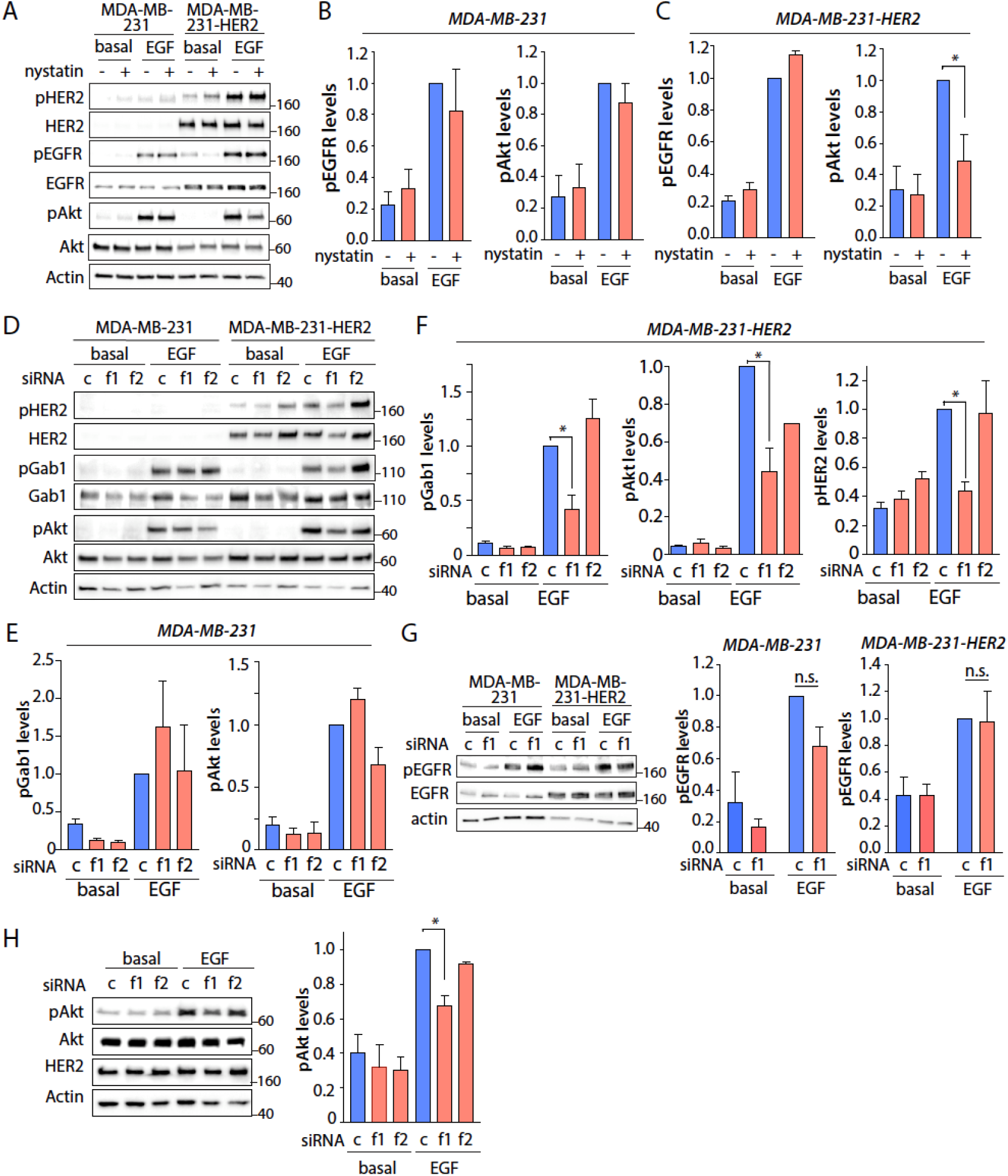
Nystatin treatment or flotillin-1 silencing selectively impairs EGFR signaling in cells expressing HER2. (**A-C**) MDA-MB-231 or MDA-MB-231-HER2 cells were treated with 50 µM nystatin or vehicle (DMSO) control for 1 h and then stimulated with 5 ng/mL of EGF for 5 min. Shown in (A) are representative western blots of whole cell lysates with antibodies as indicated and in (B-C) the levels of each of pAkt and pEGFR relative to corresponding total protein levels, as the mean ± SE; n = 3 independent experiments; *, p < 0.05 (**D-G**) MDA-MB-231 or MDA-MB-231-HER2 cells were subjected to flotillin-1 (f1) or flotillin-2 (f2) silencing, or treated with non-targeting control siRNA (c) and then stimulated with 5 ng/mL of EGF for 5 min. Shown in (D) are representative western blots of whole cell lysates with antibodies as indicated, and in (E-G) the levels of each of pGab1, pAkt, pHER2 or pEGFR as indicated, relative to corresponding total protein levels, as mean ± SE; n > 3 independent experiments *, p < 0.05. (**H**) SK-BR-3 cells were subjected to flotillin-1 (f1) or flotillin-2 (f2) silencing, or treated with non-targeting control siRNA (c) and then stimulated with 5 ng/mL of EGF for 5 min. Shown (left panel) are representative western blots of whole cell lysates with antibodies as indicated and (right panels) the levels of pAkt, relative to total Akt levels, as mean ± SE; n = 3 independent experiments. *, p< 0.05. All western blot panels show approximate molecular weight in kDa.

Membrane rafts are a heterogeneous collection of dynamic membrane structures that includes flotillin- and caveolin-based structures. To determine if the effect of nystatin treatment in MDA-MB-231-HER2 cells was due to a requirement for flotillin nanodomains, we examined the effect of flotillin-1 or flotillin-2 silencing, which we had established previously (Fekri et al., 2019). Silencing of flotillin-1 impaired EGF-stimulated Gab1 and Akt phosphorylation selectively in MDA-MB-231-HER2 cells (**Figure 2D, F**) but not in parental MDA-MB-231 cells (**Figure 2D-E**). In contrast, silencing of flotillin-2 was without effect on signaling in either cell type. That flotillin-1 silencing exhibits distinct effects from that of flotillin-2 in EGF stimulated signaling is consistent with previous findings of distinct, non-redundant roles of flotillin1 and 2 (Bitsikas et al., 2014; Kurrle et al., 2012; Otto and Nichols, 2011). This indicates that flotillin-1 is essential for EGF-stimulated activation of the PI3K-Akt pathway in MDA-MB-231 cells co-expressing HER2 but not parental cells that lack detectable HER2 expression.

Interestingly, silencing of flotillin-1 impaired EGF-stimulated HER2 phosphorylation in MDA-MB-231-HER2 cells (**Figure 2D, F**). In contrast, flotillin-1 silencing was without effect on EGF-stimulated EGFR phosphorylation in either cell line (**Figure 2G**). This suggests that similarly to the effects of nystatin, flotillin-1 silencing does not impact EGFR cell surface expression levels or ligand binding. Taken together with the altered EGF-stimulated signaling upon co-expression of HER2 indicating the formation of EGFR-HER heterodimers, these experiments indicate that cells co-expressing HER2 require flotillin-1 for multiple facets of signaling, including phosphorylation of HER2 within an EGFR/HER2 heterodimer and PI3K-Akt signaling downstream of the receptor.

To exclude the possibility that the effects of HER2 overexpression on EGFR signaling were unique to MDA-MB-231 cells, we examined ARPE-19 cells, which also lack endogenous HER2 expression. We previously similarly engineered a derivative of these cells that stably expresses HER2 (ARPE-19-HER2) (Garay et al., 2015). Consistent with our results in MBA-MB-231 cells, ARPE-19 cells stably expressing HER2 exhibit loss of EGF-stimulated Akt phosphorylation upon flotillin silencing, while parental ARPE-19 cells that lack HER2 were unaffected by flotillin silencing (**Figure S1**). Furthermore, silencing of flotillin-1 impaired EGF-stimulated Akt phosphorylation in Sk-Br-3 cells, a breast cancer cell line that endogenously expresses EGFR and high levels of HER2 (**Figure 2H**). Collectively, these results indicate that cholesterol and flotillins have a selective role in promoting EGFR signaling in cells that express HER2, but not in cells that lack HER2 expression. This in turn suggests that EGFR and/or its signals may be selectively localized and/scaffolded in flotillin nanodomains upon HER2 co-expression.

### EGFR localizes to flotillin nanodomains selectively upon HER2 co-expression

To determine how HER2 co-expression may alter the spatial organization of EGFR with respect to flotillin nanodomains and clathrin coated pits at the plasma membrane, we used total internal reflection fluorescence microscopy (TIRF-M) and labeling of the receptor and markers for each plasma membrane nanodomain. Labeling of ligand-bound EGFR was achieved by treatment with 20 ng/mL rhodamine-EGF (rho-EGF), while labeling of flotillin and clathrin nanodomains was achieved by immunofluorescence staining using specific antibodies for flotillin-1 (validated previously, (Fekri et al., 2019)) and the clathrin adaptor protein AP2 (**Figure 3A**). We also labeled cells with fluorescently conjugated trastuzumab to confirm expression of HER2. MDA-MB-231-HER2 cells exhibit some flotillin-1 structures that were also positive for rho-EGF, while these structures could not readily be observed in MDA-MB-231 parental cells that lacked HER2 expression (**Figure 3A**).

**Figure 3.**
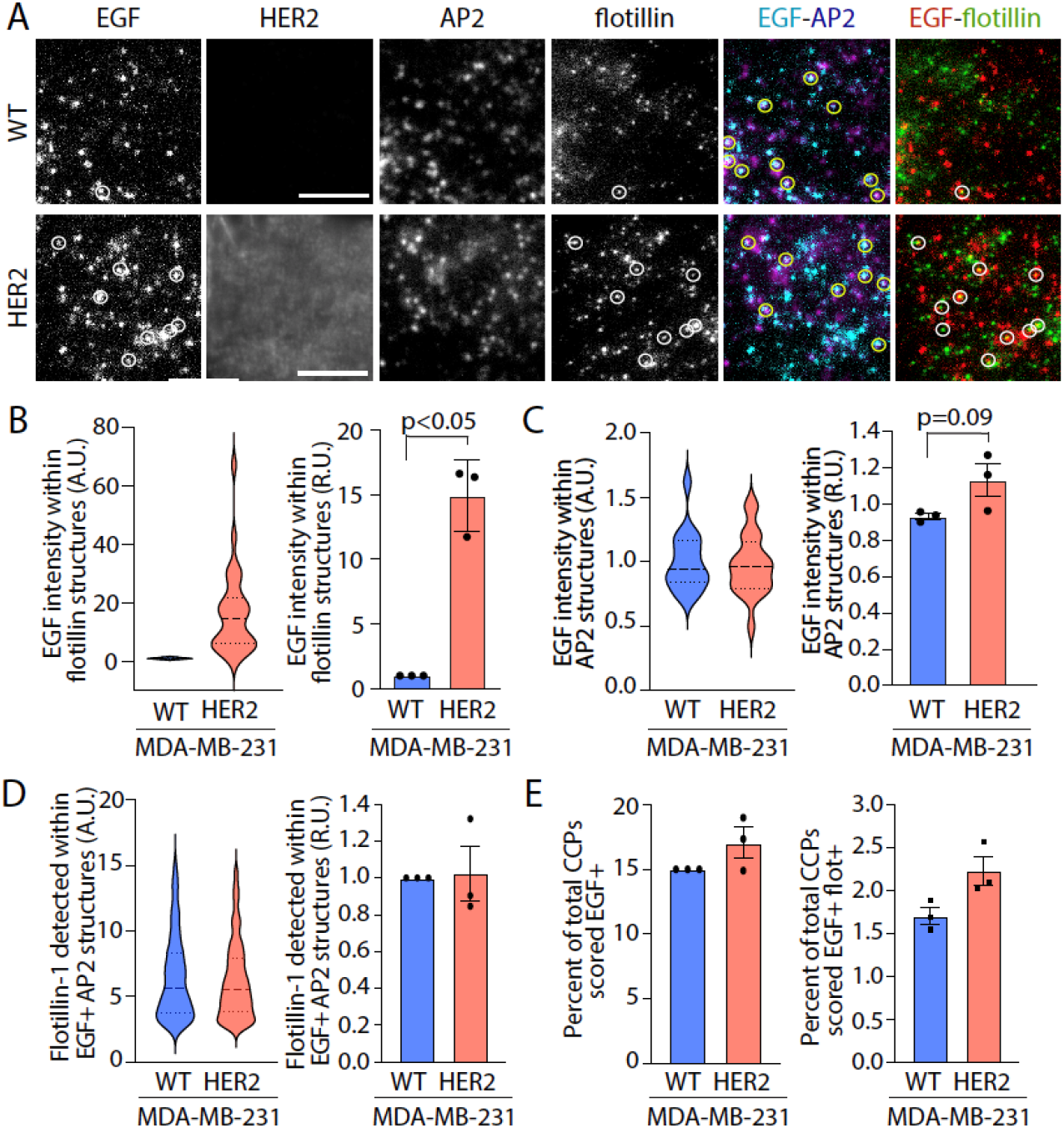
Ligand-bound EGFR is selectively recruited to flotillin structures in HER2-expressing cells. MDA-MB-231 and MDA-MB-231-HER2 cells were treated with 20 ng/mL rhodamine-EGF, fixed and labelled with flotillin-1 and AP2 antibodies. (**A**) Shown are representative images obtained by TIRF-M; scale 3 µm, (**B**) Images were subjected to automated detection of flotillin-1 structures followed by the quantification of the mean fluorescence intensity of rho-EGF therein; shown are measurements of mean rho-EGF intensity in flotillin-1 objects in individual cells (left panel, violin plot) and the mean of independent experiments (right panel). (**C-D**) Images obtained were subjected to automated detection of AP2 (clathrin) structures (clathrin-coated pits, CCPs) followed by the quantification of the mean fluorescence intensity of rho-EGF (C) or flotillin-1 (D) therein; shown for each are measurements of mean rho-EGF/flotillin intensity in AP2 (clathrin) objects in individual cells (left panel, violin plot) and the mean of independent experiments (right panel). (**E**) Following detection of AP2 (clathrin) objects as in C-D, individual AP2 objects (CCPs) were sorted based on EGF and flotillin-1 content as described in *Materials and Methods*. Shown are the percent of total AP2 structures (CCPs) that are EGF+ or EGF+/flotillin+ in each cell line, as the mean (bars) ± SE of three independent experiments (dots). The total number of flotillin structures, AP2 structures, and cells (respectively) analyzed in n = 3 independent experiments were as follows: MDA-MB-231: 27516, 53951, 48; MDA-MB-231-HER2: 33570, 64596, 79.

To quantitatively compare rho-EGF localization in flotillin structures in MDA-MB-231 parental and HER2 cells, we used automated detection and analysis of these diffraction-limited objects using a Gaussian-based modeling approach (Aguet et al., 2013), as we have used previously to detect signaling proteins within various plasma membrane structures (Cabral-Dias et al., 2022; Delos Santos et al., 2017; Lucarelli et al., 2016). We first used this method for detection of flotillin-1 objects. Using this method, we found that rho-EGF was virtually undetectable in flotillin-1 structures in parental MDA-MB-231 cells that lack HER2. In contrast, MDA-MB-231-HER2 cells had a ∼15-fold increase in rho-EGF enrichment in flotillin structures, observed when examining the levels of rho-EGF in flotillin structures in individual cells within an experiment (**Figure 3B**, *left panel*), and the mean rho-EGF within flotillin structures in independent experiments (**Figure 3B**, *right panel*). Since rho-EGF binds directly to EGFR and not to HER2, this indicates that HER2 co-expression alters the distribution of EGFR to cause enrichment of EGFR within flotillin nanodomains, likely as part of an EGFR/HER2 heterodimer.

EGF binding triggers recruitment of EGFR to clathrin-coated pits (CCPs) at the plasma membrane, which eventually leads to receptor endocytosis. The increased recruitment of EGFR to flotillin-1 structures in HER2 expressing cells could indicate that in HER2-expressing cells, there may be either loss of EGFR recruitment to plasma membrane clathrin structures, reflecting competitive recruitment of a limited pool of EGFR to each nanodomain, or recruitment of EGFR to both types of structure. To resolve this, we used primary detection of clathrin structures via AP2 labeling similarly as we had done for flotillin-1 structures. This revealed a comparable level of rho-EGF detected within CCPs in MDA-MB-231-HER2 cells as parental cells (**Figure 3C**). This indicates that in HER2-expressing cells, EGFR is recruited to clathrin-coated pits that likely support the internalization of EGFR/HER2 heterodimers as previously reported, while a pool of EGFR/HER2 heterodimers is also recruited to flotillin nanodomains. As HER2 expression was significantly more than that of endogenous EGFR (**Figure 1C**), it is likely that the majority of ligand-bound EGFR participates in EGFR/HER2 heterodimers, such that the localization of rho-EGF to flotillin and clathrin objects largely reflects that of EGFR/HER2 heterodimers.

That EGFR is recruited to both clathrin and flotillin plasma membrane structures in HER2 expressing cells may suggest that separate pools of EGFR/HER2 heterodimers are recruited to each type of structure, either sequentially or in parallel. Alternatively, HER2 expression may lead to the recruitment of flotillins to clathrin-coated pits, such that EGFR/HER2 heterodimers reside in structures comprised of both clathrin and flotillins that are not present in cells lacking HER2 expression. As such, we aimed to resolve whether HER2 co-expression may lead to alterations in the overlap of flotillin and clathrin objects at the cell surface. To do so, we examined the levels of flotillin-1 within clathrin objects detected via AP2 labeling. The median level of flotillin-1 within individual clathrin objects in a typical experiment was negative, both in MDA-MB-231 parental cells as well as in MDA-MB-231-HER2 cells (**Figure S2**). As the levels of each protein in each object is determined by the amplitude of a Gaussian modeling at the object position, negative flotillin-1 levels represent a tendency towards mutual exclusivity of flotillin-1 and clathrin. However, there was nonetheless a portion of the distribution of flotillin-1 levels within clathrin objects that could reflect some overlap of a subset of clathrin objects and flotillin (**Figure S2**).

We reasoned that if there was a minor subset of clathrin coated pits that exhibited overlap with flotillin in cells expressing HER2, this would be the subset of clathrin structures positive for EGFR/HER2 heterodimers, identified by rho-EGF detection. To probe if this is the case, we used an arbitrary but systematic threshold to score clathrin structures as EGF+ as done previously (Cabral-Dias et al., 2022; Delos Santos et al., 2017). This revealed that EGF+ clathrin structures exhibited indistinguishable flotillin levels in parental MDA-MB-231 cells and in MDA-MB-231-HER2 cells, both in individual cells within an experiment (**Figure 3D**, *left panel*), and the mean rho-EGF within flotillin structures in independent experiments (**Figure 3D**, *right panel*). We also similarly used the 5^th^ percentile of the intensity of flotillin objects determined by detection of flotillin as the primary channel to sort detected clathrin objects into flotillin+ and flotillin-cohorts. Using these sorting approaches, we found indistinguishable fractions of EGF+ clathrin objects (**Figure 3E**, *left panel*) and EGF+ flotillin+ clathrin objects (**Figure 3E**, *right panel*) in MDA-MB-231 parental cells and MDA-MB-231-HER2 cells.

Collectively, these experiments indicate that the distinct requirement for flotillin-1 in MDA-MB-231 cells that co-express HER2 correlates with an increased recruitment of EGFR (and thus likely EGFR/HER2 heterodimers) to flotillin structures, compared to cells lacking HER2 expression. These results also suggest that a subset of EGFR/HER2 heterodimers are recruited to flotillin structures that are distinct from clathrin structures, such that in HER2-expressing cells flotillin is required for activation of PI3K-Akt signaling, a major regulator of cell proliferation and survival.

### Flotillin is selectively required for cell proliferation in HER2-expressing cells

To determine how the alteration in EGFR spatial organization to flotillin nanodomains and flotillin requirement of EGFR signaling upon HER2 co-expression broadly impacts cell physiology, we next examined the effect of flotillin silencing on cell proliferation and survival. To do so, we used an automated imaging system and quantified cellular confluence in phase-contrast images over time. MDA-MB-231 (parental) cells exhibited a time-dependent increase in cell confluence indicating cellular proliferation, which was unaffected by flotillin-1 silencing (**Figure 4A**). In contrast, the increase in cell confluence reflecting proliferation in MDA-MB-231-HER2 cells was significantly impaired by flotillin-1 silencing (**Figure 4B**). Collectively, these results indicate that HER2 co-expression leads to a shift in the spatial organization of EGFR, such that the receptor becomes enriched in flotillin nanodomains, and that this localization of EGFR/HER2 heterodimers to flotillin nanodomains is required to support EGFR signaling, and cell proliferation and/or survival.

**Figure 4.**
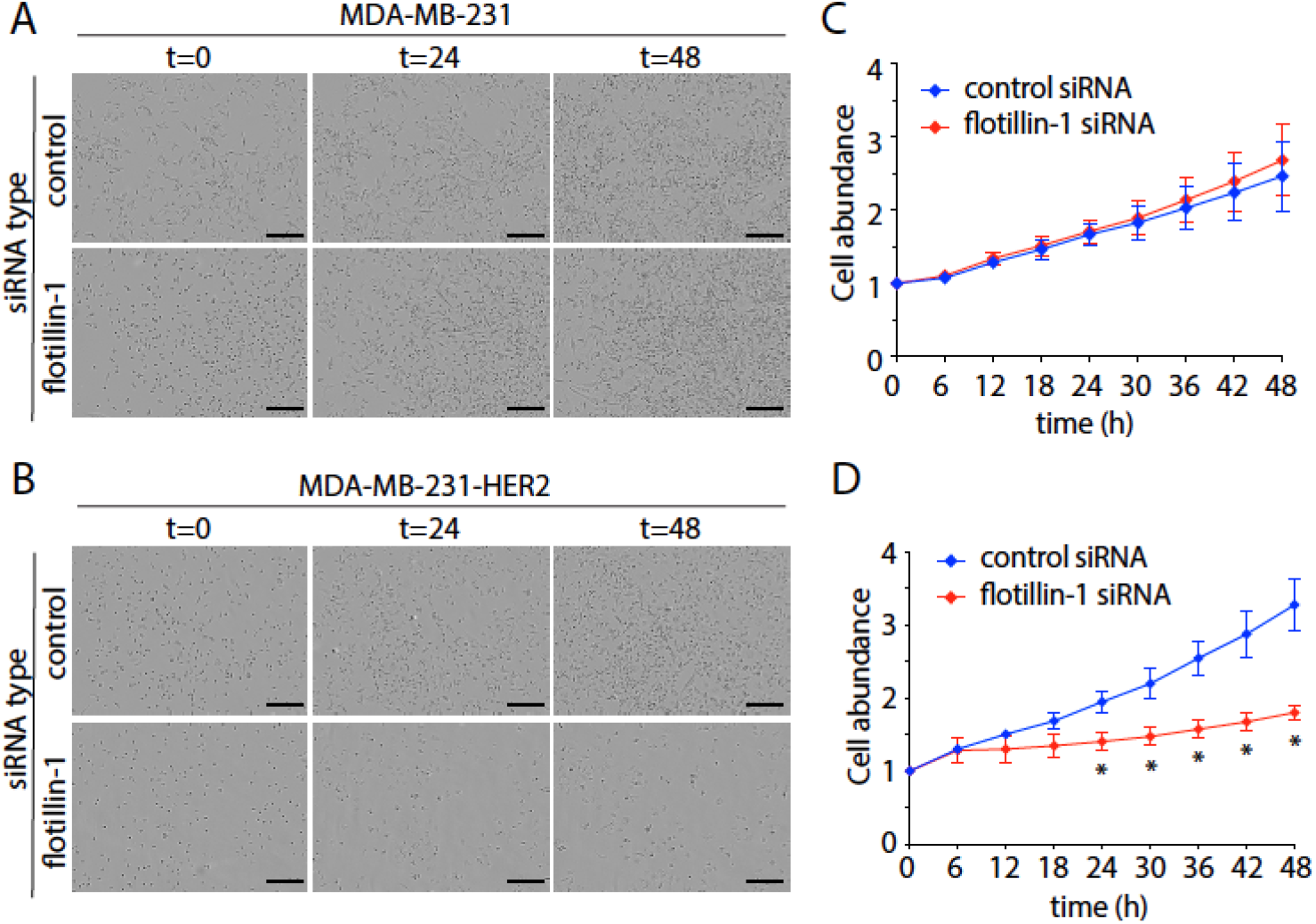
Flotillin-1 silencing selectively impairs cellular proliferation and/or survival in cells expressing HER2. MDA-MB-231 and MDA-MB-231-HER2 cells were subjected to flotillin-1 silencing or treatment with non-targeting (control) siRNA. Following transfection, cells were subject to time-lapse imaging in an IncuCyte SX5 incubated microscope for 48 h to measure cellular growth over time. (**A-B**) Shown are representative phase contrast images of MDA-MB-231 (A) and MDA-MB-231-HER2 (B) cells Scale, 250 µM. (**C-D**) Quantification of cellular confluence measured from the phase contrast images, normalized to t = 0 h, control siRNA condition, shown as the mean ± SE of n = 3 independent experiments in MDA-MB-231 (C) and MDA-MB-231-HER2 (D) cells. *, p< 0.05;

### c-Src is selectively required for EGFR signaling upon HER2 co-expression

The unique requirement for flotillin-1 for EGFR signaling and recruitment of the receptor to plasma membrane flotillin structures in HER2 expressing cells suggests that HER2 expression may also induce a unique requirement for specific signaling intermediates that may be scaffolded within flotillin nanodomains. A prime candidate for such a unique signaling intermediate is c-Src, given that this kinase is able to interact directly with HER2 but not EGFR (Kim et al., 2005; Marcotte et al., 2009). Thus, we next sought to determine if c-Src may be uniquely required for EGFR signaling upon HER2 co-expression. Silencing of c-Src (**Figure S3**) impaired EGF-stimulated Akt phosphorylation in MDA-MB-231-HER2 cells (**Figure 5B**) but was without effect in parental MDA-MB-231 cells that lacked HER2 expression (**Figure 5A**). Furthermore, and consistent with the effects of flotillin-1 silencing, c-Src silencing led to a reduction of EGF-stimulated HER2 phosphorylation (**Figure 5C**), but not of EGF-stimulated EGFR phosphorylation (**Figure 5D**). The similarity of the phenotypes observed upon flotillin and c-Src silencing on signaling by EGFR, specific for HER2 expressing cells, suggests a common mechanism of action of c-Src and flotillin in the regulation of signaling upon EGF stimulation.

**Figure 5.**
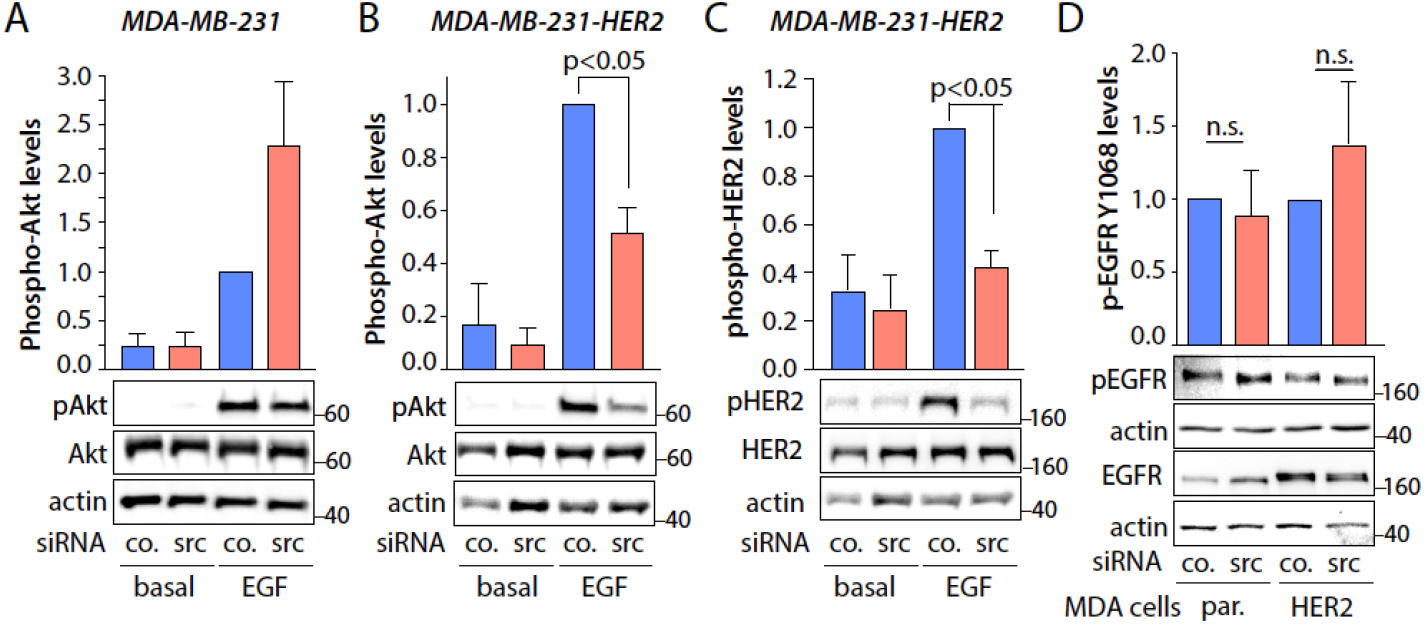
c-Src silencing selectively impairs EGFR signaling in HER2 expressing cells. MDA-MB-231 and MDA-MB-231-HER2 cells were subjected to c-Src silencing or treatment with control (non-targeting) siRNA and then stimulated with 5 ng/mL EGF for 5 min. Shown (*bottom panels*) are representative western blots of whole cell lysates probed with antibodies detecting pAkt in MDA-MB-231 (A) or in MDA-MB-231-HER2 cells (B), or pHER2 in MDA-MB-231-HER2 cells (C), or pEGFR levels in EGF-stimulated MDA-MB-231 and MDA-MB-231-HER2 (D). The levels of each of pAkt, pHER2 and pEGFR are also shown (*top panels*) relative to corresponding total protein levels, as mean ± SE of n = 3 independent experiments. All western blot panels show approximate molecular weight in kDa.

c-Src can interact directly with HER2 via binding to the HER2 kinase domain with a motif surrounding Y877 on HER2 (Kim et al., 2005; Marcotte et al., 2009). Flotillin nanodomains are enriched in gangliosides and Gb3 co-immunoprecipitates with c-Src, suggesting formation of a transbilayer spanning complex (Roy et al., 2020). This may indicate that flotillin nanodomains may contribute to the selective engagement of c-Src by EGFR/HER2 heterodimers. To test this possibility, we first examined if c-Src was recruited to flotillin nanodomains. We transfected cells with eGFP-tagged c-Src (cSrc-eGFP), following by imaging using TIRF-M and automated image analysis that allowed measurement of c-Src fluorescence intensity within flotillin structures (**Figure 6A-B**). c-Src fluorescence intensity was increased significantly in flotillin-1 structures by EGF stimulation in MDA-MB-231-HER2 cells (**Figure 6B**, *right panels*), while EGF stimulation did not impact c-Src levels in flotillin structures in parental MDA-MB-231 cells lacking HER2 expression (**Figure 6B**, *left panels*). As shown in earlier figures, in **Figure 6** we present measurements of the levels of c-Src in flotillin objects in individual cells within an experiment (violin plots) as well as the measurements from several independent experiments (bar graphs with individual data points). This indicates that HER2 expression alongside EGF stimulation contributes to the recruitment of c-Src to flotillin structures.

**Figure 6.**
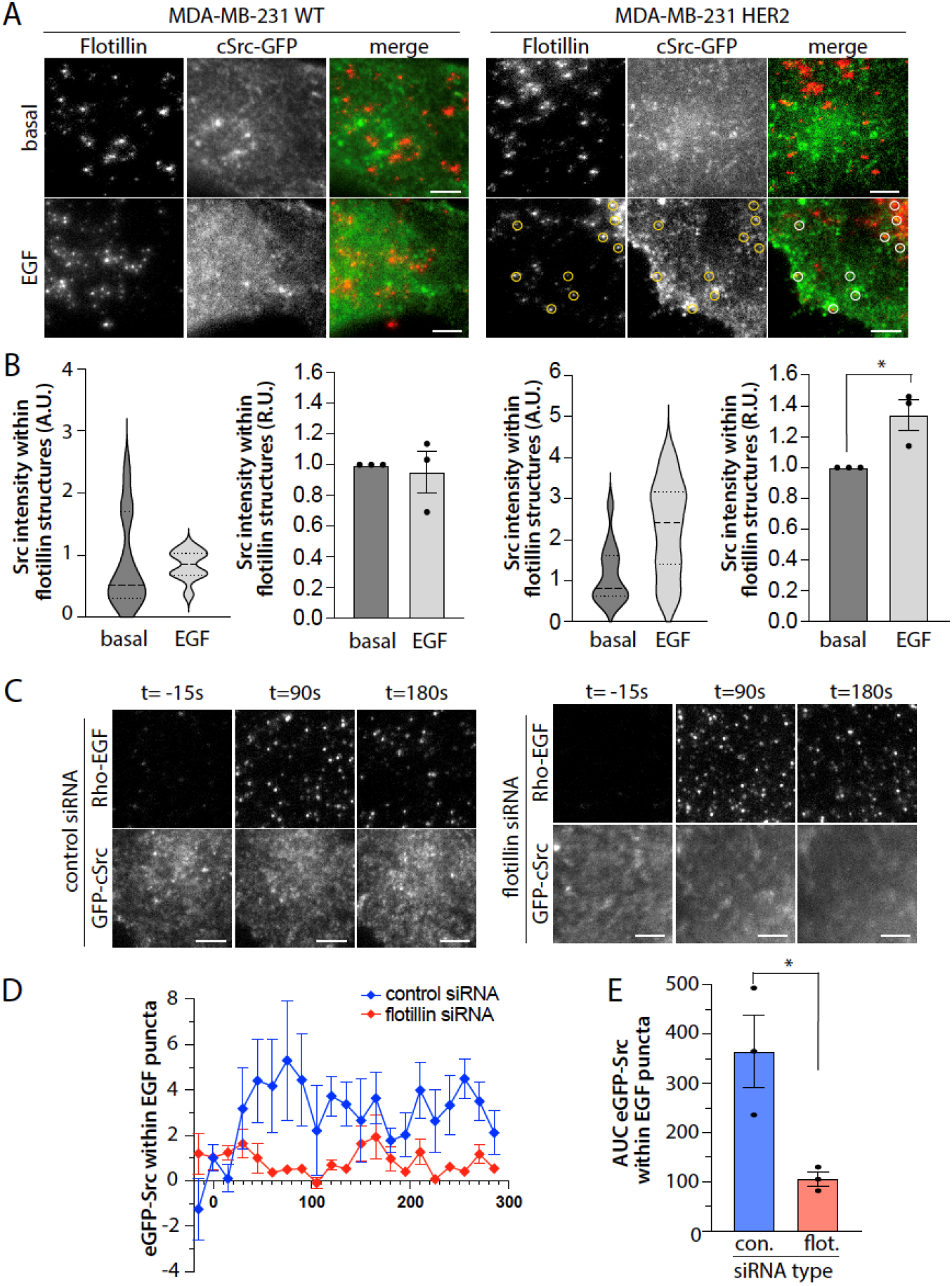
c-Src is selectively recruited to flotillin structures in cells expressing HER2. (**A-B**) MDA-MB-231 and MDA-MB-231-HER2 cells were subjected to flotillin-1 silencing or treatment with non-targeting (control) siRNA, and c-Src-eGFP transfection, stimulated with 20 ng/mL EGF for 5 min, then fixed and labelled with flotillin-1 antibodies. (A) Shown are the representative images obtained by TIRF-M; scale bar: 5 µm. (B) Images obtained were subjected to automated detection of flotillin-1 structures followed by the quantification of the mean fluorescence intensity c-Src-eGFP therein; shown are measurements of mean c-Src-eGFP intensity in flotillin-1 objects in individual cells (left panel, violin plot) and the mean of independent experiments (right panel), for each of MDA-MB-231 and MDA-MB-231-HER2 as indicated. The total number of flotillin-1 structures and cells (respectively) analyzed in n = 3 independent experiments were as follows: MDA-MB-231 WT basal: 11873, 40; MDA-MB-231 WT EGF: 15216, 39; MDA-MB-231-HER2 basal: 14282, 49; MDA-MB-231-HER2 EGF: 17151, 54. (**C-E**) MDA-MB-231-HER2 cells were subjected to flotillin-1 silencing or treatment with non-targeting (control) siRNA, and c-Src-eGFP transfection. Cells were then subjected to live-cell TIRF-M; stimulation with 20 ng/mL of rhodamine-EGF begins at t = 0 min followed by imaging for an additional 285s at 15 frames per second. Shown in (C) are representative images obtained by TIRF-M; scale bar = 5 µM. Images obtained were subjected to automated detection of rho-EGF structures followed by the quantification of the mean fluorescence intensity of c-Src-eGFP therein; shown in (D) are measurements of the mean ± SE c-Src-EGFP intensity in rho-EGF objects from independent cell measurements, normalized to the mean c-Src-eGFP fluorescence in rho-EGF objects at time 0-30 s of observation. >50 objects per frame were analyzed from >7 cells in 3 independent experiments. Shown in (E) are area under the curve (AUC) measurements for each independent experiment from the data in (D), * p < 0.05.

This suggests that while HER2 may interact with c-Src directly, the interaction of c-Src with HER2 may further require flotillin nanodomains, perhaps as a form of coincidence detection. To test this, we examined the colocalization of GFP-tagged c-Src with EGFR (labeled with rho-EGF) in MDA-MB-231-HER2 cells, alongside flotillin-1 silencing using time-lapse TIRF-M (**Figure 6C**). We used automated detection of rho-EGF puncta by Gaussian modeling of the point spread function of these diffraction-limited objects, followed by quantification of c-Src-eGFP fluorescence within these EGFR objects. These experiments reveal that EGF stimulation causes a rapid recruitment of c-Src-eGFP to EGFR objects in MDA-MB-231-HER2 cells treated with control (non-targeting) siRNA (**Figure 6D**). In contrast, MDA-MB-231-HER2 cells treated with flotillin-1 siRNA exhibited no detectable EGF-stimulated increase in c-Src-eGFP within EGFR objects (**Figure 6D**). Quantification of the area under the curve for these time-lapse measurements of c-Src-GFP recruitment to EGFR puncta showed a significant reduction of this phenomenon in flotillin-1 silenced cells (**Figure 6E**). Taken together, these results suggest that flotillin-1 serves as a part of a coincidence detection mechanism alongside HER2 to recruit c-Src to EGFR/HER2 signaling complexes, underlying a unique functional requirement for c-Src in signaling by EGFR/HER2 heterodimers.

Since we observed a unique role for flotillin nanodomains in controlling cell proliferation (**Figure 4**), we next sought to determine if c-Src may also similarly be required for control of cell proliferation selectively in HER2 expressing cells. As we had observed in Figure 4A, MDA-MB-231 (parental) cells exhibited a time-dependent increase in cell confluence indicating cellular proliferation, which was unaffected by c-Src silencing (**Figure 7A**). In contrast, the increase in cell confluence reflecting proliferation in MDA-MB-231-HER2 cells was significantly impaired by c-Src silencing (**Figure 7B**). These results suggest that the unique requirement for c-Src in MDA-MB-231 HER2 cells but not parental MDA-MB-231 cells lacking HER2 for EGFR signaling is mirrored by a unique requirement in HER2-expressing cells for c-Src for cell proliferation.

**Figure 7.**
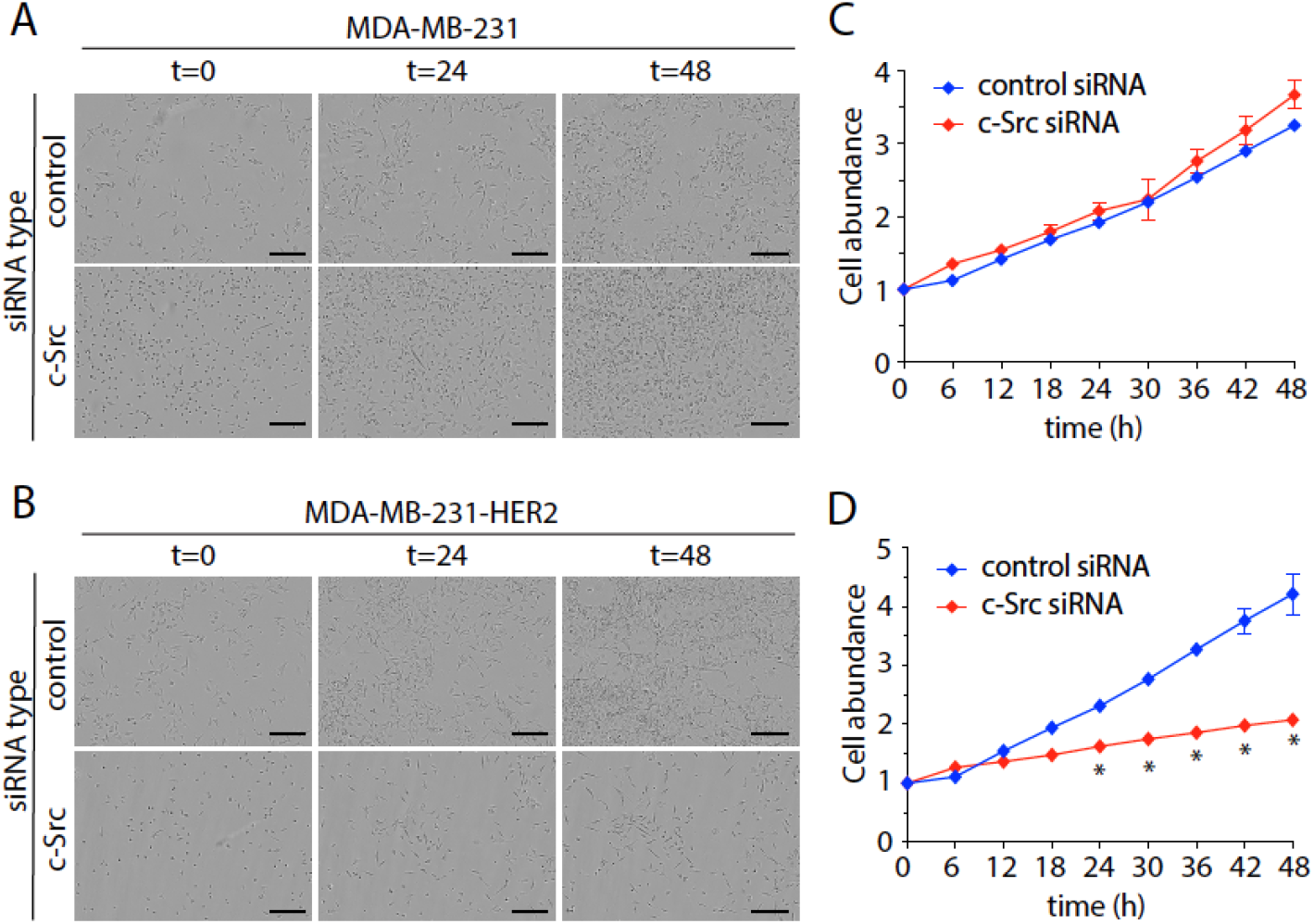
c-Src silencing selectively impairs cellular proliferation and/or survival in cells expressing HER2. MDA-MB-231 and MDA-MB-231-HER2 cells were subjected to c-Src silencing or treatment with non-targeting (control) siRNA. Following transfection, cells were subject to time-lapse imaging in an IncuCyte SX5 incubated microscope for 48 h to measure cellular growth over time. (**A-B**) Shown are representative phase contrast images of MDA-MB-231 (A) and MDA-MB-231-HER2 (B) cells Scale, 250 µM. (**C-D**) Quantification of cellular confluence measured from the phase contrast images, normalized to t = 0 h, control siRNA condition, shown as the mean ± SE of 3 independent experiments in MDA-MB-231 (C) and MDA-MB-231-HER2 (D) cells. *, p< 0.05;

## Discussion

We found a selective requirement for flotillin and c-Src in EGF-stimulated activation of EGFR signaling leading to Akt phosphorylation in cells expressing HER2. Consistent with this, in cells co-expressing HER2, EGFR exhibited increased localization to plasma membrane flotillin structures. Cell proliferation and/or survival was selectively dependent on flotillin and c-Src in cells co-expressing HER2. Hence, we propose that HER2 co-expression defines a shift in EGFR signaling to a flotillin membrane raft- and c-Src-dependent mechanism.

### Regulation of EGFR signaling by membrane rafts

Many studies have examined whether and how EGFR may be localized to membrane raft domains. Some studies relied on isolation of raft-enriched membrane fractions (reviewed by (Delos Santos et al., 2015; Lu and Fairn, 2018)). Other methods have relied on co-localization of EGFR with membrane raft markers such as cholera toxin or GM1 based on fluorescence (Hofman et al., 2008) or electron (Puri et al., 2005) microscopy methods. The effect of cholesterol perturbation on EGFR mobility can also be measured by single particle tracking, which allows inference of the localization of EGFR to cholesterol-dependent membrane rafts (Michio Hiroshima et al., 2021; Orr et al., 2005). Collectively, these studies provide evidence that EGFR resides within dynamic membrane raft domains, at least under some cellular contexts.

In addition to examining membrane raft localization, some previous studies also investigated the role of membrane rafts in EGFR signaling, largely by cholesterol disruption. These can be broadly classified into three categories based on their findings: 1) that cholesterol disruption increases EGFR ligand binding and/or signaling (Chen and Resh, 2002; Lambert et al., 2008; Li et al., 2006; Liu et al., 2009; Michio Hiroshima et al., 2021; Pike and Casey, 2002; Ringerike et al., 2002; Roepstorff et al., 2002; Turk et al., 2012; Westover et al., 2003; Zhang et al., 2016)., 2) that cholesterol disruption has little or no effect on EGFR signaling (Sigismund et al., 2008; Ushio-Fukai et al., 2001; Zhang et al., 2016) and 3) that cholesterol disruption impairs EGFR signaling (Gibson et al., 2009; Hea et al., 2007; Irwin et al., 2011).

Since these studies suggest that EGFR distinctly requires cholesterol and membrane rafts in some cellular contexts, the expression of other specific protein(s) may shift EGFR nanoscale organization to membrane rafts, leading to altered requirements for signaling. Our results suggest that the availability of HER2 to form heterodimers with EGFR is one of the factors that defines this switch in regulation by cholesterol and membrane rafts. However, few of these prior studies directly examined the relative expression of EGFR and HER2 or examined the effect of alterations in HER2 expression in a common cellular background, as we have done here. Our results show that HER2 co-expression induces cholesterol- and flotillin-dependence for EGFR signaling and EGFR localization to flotillin nanodomains.

### Regulation of EGFR/HER2 signaling by flotillin nanodomains

Flotillin-1 and -2 each contain an N-terminal prohibitin homology (PHB) domain also known as a SPFH (stomatin/prohibitin/flotillin/HflK/C) domain. This PHB domain has membrane insertion domains, S-acylation sites and mediates interaction with cholesterol (Babuke et al., 2009; Kurrle et al., 2012; Li et al., 2012a; Meister and Tikkanen, 2014; Morrow et al., 2002; Neumann-Giesen et al., 2004; Staubach and Hanisch, 2011; Strauss et al., 2010). An additional C-terminal domain mediates oligomerization (Babuke et al., 2009; Grant & Benjamin, 2011; Solis et al., 2007). These properties allow flotillins to form membrane nanodomains that are enriched in cholesterol, glycosphingolipids, as well as some signaling proteins.

Signaling by activated dimers involving EGFR requires formation of an asymmetric kinase dimer, in which one kinase domain serves as the “activator”, and the other as the “receiver”, such that only the receiver kinase is active (Jura et al., 2009; Zhang et al., 2006). Importantly, in the context of EGFR/HER2 heterodimers, the HER2 kinase preferentially adopts the activator (inactive) position (Macdonald-Obermann et al., 2012; Ward and Leahy, 2015). Thus, phosphorylation of HER2 upon EGF stimulation is expected to result from direct phosphorylation by the EGFR kinase domain or via an indirect mechanism, such as by c-Src or other kinase.

We observe that flotillin-1 and c-Src are required for EGF-stimulated phosphorylation of Akt and Y1248 on HER2, but not the phosphorylation of EGFR Y1068. This indicates that perturbations of flotillin-1 or c-Src do not alter EGFR cell surface levels or ligand binding, but instead may alter signaling by EGFR downstream of EGFR/HER2 heterodimerization. One possibility is that cholesterol and/or other lipids in flotillin membrane rafts impacts the conformation of the EGFR/HER2 heterodimer, gating phosphorylation of certain receptor residues such as Y1248 on HER2. Interestingly, computational modeling indicates that the HER2 transmembrane domain may bind cholesterol, impacting the preference for the formation of several possible dimer configurations by the HER2 transmembrane domain (Pawar and Sengupta, 2021). This suggests that cholesterol present within flotillin nanodomains may uniquely regulate the conformation of EGFR/HER2 heterodimers to favor EGF-stimulated phosphorylation of Y1248 and other residues essential for signal transduction.

In addition, flotillin nanodomains can also regulate EGFR signaling by enrichment of signaling regulators or intermediates therein. The ganglioside Gb3 is enriched within flotillin nanodomains (Brandel et al., 2021) and despite being on opposite leaflets, Gb3 can co-immunoprecipitate with c-Src (Roy et al., 2020), suggesting formation of transbilayer spanning complexes. Indeed various Src-family kinases are associated with flotillins and flotillin nanodomains (Stuermer, 2010). Further, c-Src directly binds to the HER2 kinase domain (Kim et al., 2005; Marcotte et al., 2009). We find selective recruitment of c-Src upon EGF stimulation to plasma membrane flotillin structures which is consistent with a coincidence detection mechanism involving flotillin and HER2 for the selective recruitment of c-Src. This may in turn allow c-Src to phosphorylate specific residues on EGFR or HER2, including Y1248 on HER2. However, neither c-Src nor flotillin-1 silencing impaired EGF-stimulated phosphorylation of Y1068 on EGFR, suggesting that flotillin and c-Src are required subsequent to ligand binding and receptor heterodimerization.

Hence, we propose that the unique requirement for flotillin-1 and c-Src in signaling by EGFR/HER2 heterodimers reflects a distinct requirement for these proteins for phosphorylation of a subset of EGFR or HER2 sites and/or those of receptor-proximal signals such as Gab1 leading to Akt phosphorylation, while EGF ligand binding may be largely flotillin-1 and c-Src independent.

### Roles of flotillin versus clathrin nanodomains in EGFR/HER2 signaling

We find that HER2 expression does not suppress the recruitment of EGFR to plasma membrane clathrin structures. While some studies have reported that flotillins may be present within clathrin endocytic structures (Ge et al., 2008; Ge et al., 2011; Meister and Tikkanen, 2014), others have found these to be largely distinct (Dam et al., 2020; Fekri et al., 2019; Glebov et al., 2006). Our findings here are consistent with the latter and reveal little overlap detected between flotillin-1 and clathrin structures in MDA-MB-231 cells. Moreover, we find that HER2 expression does not alter the levels of flotillin-1 in the subset of EGF+ clathrin structures. Taken together with the observation that HER2 expression induces a robust increase in the enrichment of a subset of EGFR to flotillin nanodomains, this suggests that EGFR/HER2 heterodimers reside within either flotillin nanodomains or clathrin-coated pits, but not both simultaneously.

This in turn suggests two models of recruitment of EGFR/HER2 to clathrin and flotillin structures: sequential or parallel. In a sequential model, recruitment of EGFR/HER2 heterodimers to flotillin nanodomains precedes their transit to clathrin-coated pits, which would then lead to internalization. Plasma membrane flotillin structures are short-lived, with ∼60% of structures having lifetimes <15s (Fekri et al., 2019), which is consistent with the short residency of EGFR/HER2 in flotillin nanodomains. In the parallel model, a subset of EGFR/HER2 complexes is recruited to each nanodomain, followed perhaps by internalization therefrom. Resolving these models may require single-particle tracking of EGFR or other methods to track EGFR dynamics and are beyond the scope of this study. We focused here on how each nanodomain distinctly regulates EGF-stimulated signaling upon expression of HER2.

We and others reported that plasma membrane clathrin structures are required for regulation of EGF-stimulated Akt phosphorylation (Cabral-Dias et al., 2022; Delos Santos et al., 2017; Garay et al., 2015; Pascolutti et al., 2019; Reis et al., 2015; Rosselli-Murai et al., 2018; Sigismund et al., 2008). Specifically, we resolved that in cells that express only EGFR but not HER2, clathrin-coated pits are enriched in signaling proteins such as TOM1L1, Fyn, and SHIP2, leading to the selective activation of Akt2 (Cabral-Dias et al., 2022). Importantly, we also previously reported that expression of HER2 rendered cells insensitive to clathrin perturbation for EGF-stimulated Akt phosphorylation (Garay et al., 2015). This suggests that the signaling requirements of EGFR/HER2 heterodimers that depend on flotillin cannot be supplied by clathrin structures in the absence of flotillin, despite robust recruitment of EGFR/HER2 dimers to clathrin structures. While a subset of clathrin structures are enriched in Fyn (Cabral-Dias et al., 2022), a Src-family kinase, it is possible that the interaction of c-Src with HER2 is specific for c-Src and not Fyn, making clathrin-localized Fyn unable to compensate for loss of flotillin and/or c-Src for control of EGFR/HER2 signaling.

On the other hand, clathrin becomes dispensable for EGF-stimulated Akt phosphorylation selectively upon expression of HER2 (Garay et al., 2015). This in turn suggests that either flotillin nanodomains are redundant with the function of clathrin for activation of PI3K-Akt signaling, perhaps by scaffolding redundant signaling intermediates, or that EGFR/HER2 heterodimers do not require such nanodomain scaffolding activity once the receptor complex is fully phosphorylated. While resolving this mechanism is beyond the scope of the current study, we nonetheless establish for the first time that expression of HER2 alters the functional requirements for EGFR signaling and define cholesterol, flotillin-1, and c-Src as unique EGFR signaling intermediates in HER2 expressing cells.

Our results show that co-expression of HER2 is an important determinant of the need for flotillin nanodomains and c-Src for signal transduction and for the resulting control of cell survival and proliferation. Consistent with this, flotillin-2 expression is correlated with reduced survival and poor prognosis in HER2-positive breast cancer (Pust et al., 2013) and gastric cancer (Zhu et al., 2013). Further, HER2-positive cancers that acquire resistance to trastuzumab show elevated levels of c-Src activity, which is correlated with poor patient survival (Peiró et al., 2014; Zhang et al., 2011). While triple-negative breast cancers such as the MDA-MB-231 cells that we examine here are defined by a lack of significant expression of HER2, some reports found that a subset of these tumors may express low but detectable levels of HER2, despite no amplification of ERBB2 (Marra et al., 2020). As such, it may be useful in future studies to expand on this work to examine how even relatively low levels of expression of HER2 can influence EGFR signaling in triple negative breast cancer and other cancers.

More broadly, EGFR and HER2 are co-expressed and function in many cells and tissues, both in physiological and pathophysiological settings. Our results contribute to the understanding of cellular contexts, in this case HER2 expression, that may specify distinct requirement for membrane rafts for EGFR signaling.

## Acknowledgements

Funding for this research was provided by a Project Grant (PJT-156355) from the Canadian Institutes of Health Research (CIHR) to C.N.A. and G.D.F., and a New Investigator Award from the Canadian Institutes of Health Research (CIHR), Funding from Ryerson University, as well as an Early Researcher Award (Ontario Ministry of Research, Innovation and Science) to C.N.A. J.A was supported by an Ontario Graduate Scholarship and L.A.O. was supported by an Ontario Graduate Scholarship and a Canada Graduate Scholarship – Master’s from CIHR.

## Materials and Methods

### Materials

Antibodies used: phospho-EGFR (Y1068), phospho-HER2 (Y1248), phospho-Gab1 (Y627), flotillin, actin, c-Src, and HER2 antibodies were from Cell Signaling Technology (Danvers, MA); phospho-Akt (S473) antibody was from Invitrogen (Waltham, MA); EGFR antibody was from Santa Cruz Biotechnology (Santa Cruz, CA); ErbB2 (CD34) PE-Vio770 used for FACS sorting of HER2+ stable cells was obtained from Miltenyi Biotec (Germany).

### Cell Culture

MDA-MB-231, MDA-MB-231-HER2, ARPE-19 and SK-BR-3 cell lines were cultured as previously described (Bautista et al., 2018; Bone et al., 2017). ARPE-19 cells stably expressing HER2 were previously described (Garay et al., 2015). Cells were maintained using RPMI (Sigma Aldrich) containing 10% fetal bovine serum (Life Technologies), 100 μg/ml streptomycin and 100 U/ml penicillin (Life Technologies) at 37°C and 5% CO^2.^ MDA-MB-231-HER2 cells were grown in media supplemented with 1 ug/mL of puromycin to maintain HER2 expression.

### Generation of MDA-MB-231-HER2 stable cells

Parental MDA-MB-231 cells were transfected with pBABEpuro-ERBB2 plasmid using FuGENE HD (Promega) as per the manufacturer instructions. Briefly, cells incubated with in Opti-MEM (Life Technologies) containing reagent and plasmid at 37°C for 24 hours. HER2 positive cells were selected using media containing 3 µg/mL puromycin and then maintained in in 1 µg/mL puromycin. To obtain homogenous selection of HER2 stable cells, cell sorting was employed by labeling intact cells with the fluorescently conjugated HER2 antibody ErbB2 PE-Vio770 and then subjected to fluorescence-activated cell sorting using a FACS Aria III (BD Biosciences), selecting the cells in the top 10% of HER2 expression to generate MDA-MB-231-HER2 stable cells.

### Plasmids

pBABEpuro-ERBB2 was a gift from Matthew Meyerson (Addgene plasmid # 40978; http://n2t.net/addgene:40978; RRID:Addgene_40978). mEmerald Hc-SRC-N-18 (Src-eGFP here) was a gift from Michael Davidson (Addgene plasmid # 54118; http://n2t.net/addgene:54118; RRID:Addgene_54118)

### Plasmid and siRNA transfections

All transient transfections were performed using FuGENE HD as per manufacturer’s instructions. Cells were incubated in OptiMEM with FuGENE HD and DNA mixture at a 3:1 ratio for 4-6 h after which media containing the transfection mixture media was replaced with regular growth media. Experiments were conducted 16-18 h post transfection.

siRNA gene silencing was performed using RNAiMAX (Thermo Fisher Scientific) and customized targeted siRNA sequence as per the manufacturer’s instruction and as previously described (Delos Santos et al., 2017). Cells were seeded on a 6-well plate 24 h before transfection. Briefly, cells were initially washed with 1X PBS and replaced OptiMEM media (Thermo Fisher Scientific). RNAiMAX and the customized siRNA sequences was added to the cellular medium at a concentration of 220 pmol/L and incubated for 4-6 h before replacing transfection media with regular growth media. 2 rounds of transfection were performed, 24 h and 48 h before the experiment was conducted. The siRNA sequences were used as follows (sequence in parentheses indicates duplex overhang): control (sense): CGUACUGCUUGCGAUACGG(UU), (antisense): CCGUAUCGCAAGCAGUACG(UU) ; flotillin-1 (sense): UGGCCAAGGCACAGAGAGA(UU) (antisense): UCUCUCUGUGCCUUGGCCA(UU) ; c-Src (sense): GCAGAGAACCCGAGAGGGA(UU), (antisense): UCCCUCUCGGGUUCUCUGC(UU)

### Whole cell lysates and western blotting

This was performed as previously described (Bautista et al., 2018). Following treatment and stimulation with EGF as indicated, cells were washed with ice-cold PBS and prepared for lysis by adding 2X-Laemelli Sample buffer (0.5M Tris pH 6.8, Glycerol, 10% SDS), 1 mM sodium orthovanadate, 10 mM okadaic acid, and 20 mM of protease inhibitor, followed by passage of lysates 5x through a 27.5 gauge syringe and heating at 65°C for 15 min. 10% beta-mercaptoethanol and 5% bromophenol blue were added to the lysates prior to use of samples for SDS-PAGE. Molecular weight markers used were Novex Sharp Pre-stained Protein Standard (LC5800) or PageRuler Prestained Protein Ladder (26617) (ThermoFisher Scientific). Following SDS-PAGE, proteins were transferred to a PVDF membrane and subjected to blocking, then incubated with appropriate antibodies as previously described (Antonescu et al., 2011). Western blot signal was detected using BioRad ChemiDoc System upon soaking membranes in Luminata Crescendo HRP substrate (Millipore Sigma), ensuring that the signal was not saturated at any pixel.

Western blot images were quantified using ImageJ software (National Institutes of Health, Bethesda, MD) (Schneider et al., 2012) by signal integration in the area of each lane approximating each band. This measurement for phosphorylated proteins (e.g. pAkt) is then normalized to the loading control (e.g., actin) signal, and subsequently normalized to the total protein (e.g. Akt) signal, obtained following reblotting. In each experiment, the resulting normalized pAkt/ total Akt signal in each condition was expressed as a fraction of the normalized pAkt/total Akt measurement in the control condition stimulated with EGF for 5 min. Statistical analysis was performed by two-way (Figure 1) or one-way (Figure 2, 5) ANOVA followed by Tukey post-test, with p < 0.05 used as a threshold for establishing differences between experimental conditions.

### Cellular Inhibition and EGF stimulation

Prior to all experiments (except cell proliferation experiments), MDA-MB-231, MDA-MB-231-HER2 cells, ARPE-19, or Sk-Br-3 cells were serum starved for 1 h. For some experiments (Figure 2A-C), cells were then treated with media containing 50 µM nystatin (Sigma Aldrich, St. Louis, MI) or a corresponding volume of dimethyl sulfoxide (vehicle control) for 1 h. Unless otherwise specified, cells were stimulated with 5 ng/mL of EGF for 5 min.

### Fluorescent EGF labeling and antibody staining

Following serum starvation, cells were stimulated with 20 ng/mL rhodamine-EGF or (unlabeled) EGF for 5 min prior fixation/permeabilization with ice-cold methanol at -20°C for 20 min, as described previously to allow detection of flotillin structures (Possidonio et al., 2014). Following blocking in 3% BSA solution, cells were incubated with specific primary antibodies at room temperature for 1 h, followed by washing and incubation with appropriate secondary antibodies. Cells were maintained in PBS at 4°C prior to total internal reflection microscopy (TIRF-M) imaging.

### Total Internal Reflection Microscopy (TIRF-M) Imaging

For experiments related to Figure 3, Figures 6A,C, cell samples were imaged using a Quorum (Guelph, ON, Canada) Diskovery instrument, comprised Leica DMi8 microscope equipped with a 63×/1.49 NA TIRF objective with a 1.8× camera relay (total magnification 108×). Images from samples were acquired using a Zyla 4.2 megapixel sCMOS camera with using 488 nm, 561, or 627 nm laser illumination and 525/50, 620/60, or 700/75 emission filters. Fixed-cell TIRFM imaging (Figure 3, Figure 6A) was done at room temperature with samples mounted in PBS. For live-cell imaging experiments (Figure 6C), cells were maintained at constant 37°C during imaging, in phenol-free DMEM/F12 media (Gibco) supplemented with 20 mM HEPES and 20 ng/mL EGF, with imaging at rate of 1 frame per 15 s for 5 min.

### TIRF Microscopy Analysis to detect fluoresce intensity within AP2 (clathrin), flotillin or rho-EGF objects

Systematic, unbiased detection and analysis of either flotillin or clathrin diffraction-limited structures was done as previously described (Cabral-Dias et al., 2022; Delos Santos et al., 2017; Lucarelli et al., 2017), using custom software developed in Matlab (Mathworks Corporation, Natick, MA), as described in (Aguet et al., 2013). Briefly, diffraction limited flotillin, clathrin, or rho-EGF structures were detected using a Gaussian-based model method to approximate the point-spread function of flotillin, AP2 or EGF objects (‘primary’ channel). The TIRF-M intensity corresponding to various proteins in a ‘secondary’ (or ‘tertiary’) channel (e.g. Rho-EGF, AP2, flotillin, Src-eGFP) within each object detected in the ‘primary’ channel was determined by the amplitude of the Gaussian model for the appropriate fluorescence channel for each object position detected in the ‘primary’ channel. Sorting of objects based on flotillin-1 or rho-EGF content was done using an arbitrary yet systematic threshold in each case based on the 85^th^ percentile of EGF intensity in AP2 (clathrin) objects (Lucarelli et al., 2017), or the 5^th^ percentile of flotillin-1 intensity in flotillin-1 objects. Statistical analysis was performed by t-test, with p < 0.05 used as a threshold for establishing differences between experimental conditions.

### Cellular proliferation using IncuCyte imaging system

MDA-MB-231 and MDA-MB-231-HER2 cells were seeded on a 6 well plate and subjected to siRNA transfection as described. Immediately following the second round of transfection, cells were placed in the Incucyte SX5 and maintained at 37°C and 5% CO2 (Sartorius, Göttingen, Germany) for detection and automated analysis of cell growth/proliferation, as we described previously (Lo et al., 2022). The phase-contrast camera was used to take images every 30 min for 48 h. Time-lapse image series were analyzed using the Incucyte Basic Analysis Software. To determine cellular confluence, the phase contrast images were analyzed using the following settings: minimum area filter = 25 μm^2^: segmentation adjustment = 0.8. Measurements were normalized to initial phase contrast confluence. Statistical analysis was performed by two-way ANOVA followed by Tukey post-test, with p < 0.05 used as a threshold for establishing differences between experimental conditions.

---

**Figure S1.**
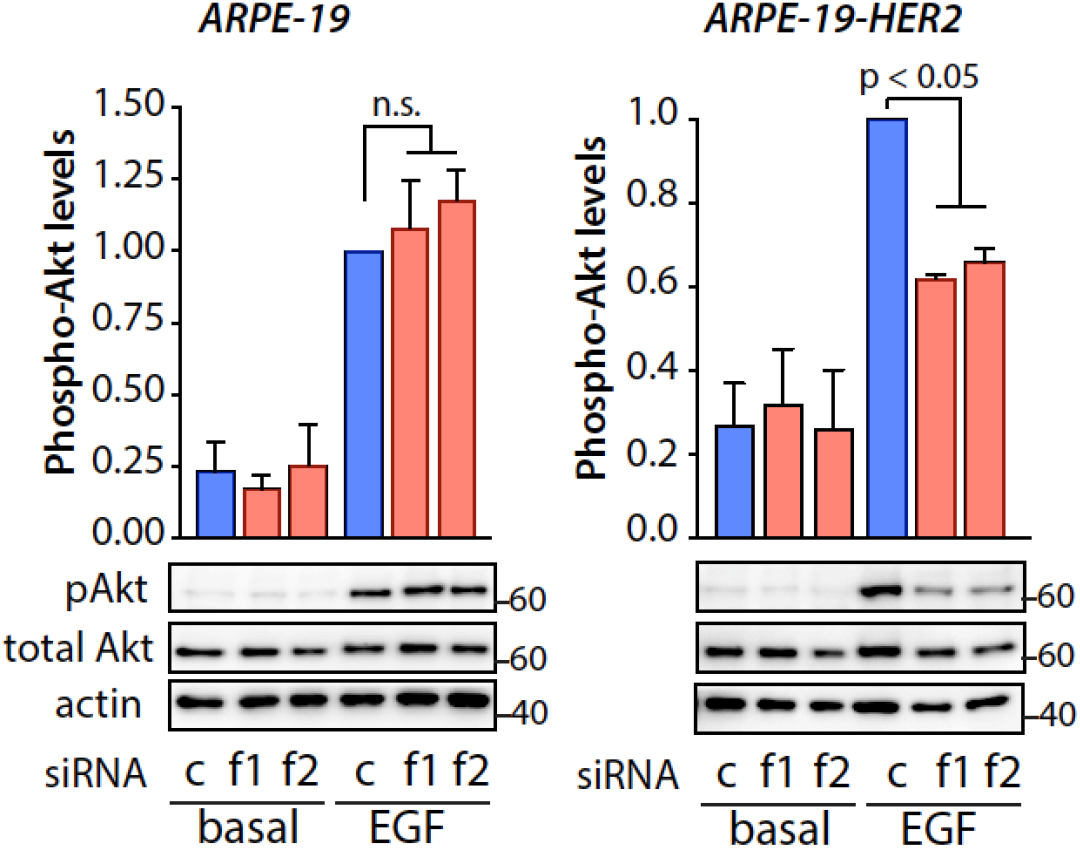
Flotillin silencing selectively impairs EGF-stimulated Akt phosphorylation in ARPE-19 cells expressing HER2. ARPE-19 cells do not express endogenous HER2 (Garay et al., 2015). We previously generated ARPE-19 cells stably expressing HER2 (ARPE-19-HER2 cells) (Garay et al., 2015). ARPE-19 or ARPE-19-HER2 cells were subjected to flotillin-1 (f1) or flotillin-2 (f2) silencing, or treated with non-targeting control siRNA (c) and then stimulated with 5 ng/mL of EGF for 5 min. Shown are representative western blots of whole cell lysates with antibodies as indicated (bottom panels), and the levels of pAkt, relative to corresponding total protein levels, as mean ± SE; n >3 independent experiments *, p< 0.05. All western blot panels show approximate molecular weight in kDa.

**Figure S2.**
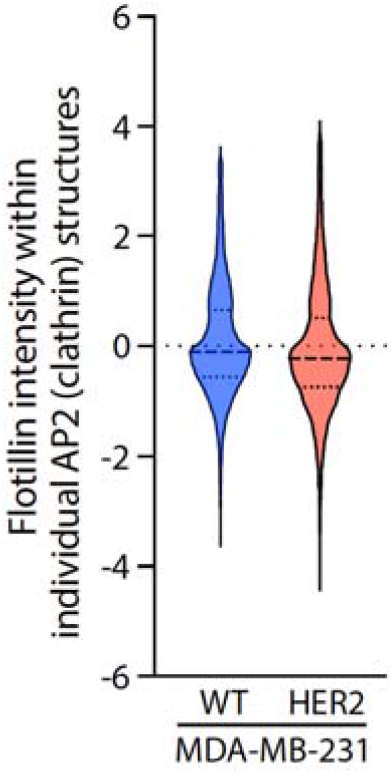
Flotillin-1 levels detected within individual AP2/clathrin plasma membrane objects. TIRF-M images obtained as described in Figure 3 were subjected to automated detection of AP2 (clathrin) structures followed by the quantification of the mean fluorescence intensity of flotillin-1 therein; shown for each are measurements of flotillin intensity in individual AP2 (clathrin) objects from a representative experiment (violin plots). The median flotillin-1 levels detected within clathrin objects is negative suggesting a tendency towards mutual exclusion. However, the distribution is not normal and the mean flotillin-1 detected within AP2 (clathrin) objects is positive, suggesting that there may be overlap of a small fraction of AP2 (clathrin) objects with flotillin. However, the analysis presented in Figure 3E indicates that expression of HER2 does not alter flotillin-1 detected within EGF+ clathrin objects.

**Figure S3.**
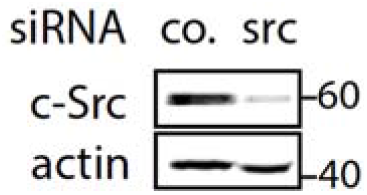
Silencing of c-Src expression. MDA-MB-231 cells were subjected to c-Src silencing (Src) or treatment with control (co; non-targeting) siRNA. Shown are representative western blots of whole cell lysates probed with antibodies detecting c-Src or actin. All western blot panels show approximate molecular weight in kDa.

